# Habitat loss exacerbates pathogen spread: An Agent-based model of avian influenza infection in migratory waterfowl

**DOI:** 10.1101/2021.10.21.465250

**Authors:** Shenglai Yin, Yanjie Xu, Mart C.M. de Jong, Mees R.S. Huisman, Andrea Contina, Herbert H. T. Prins, Zheng Y. X. Huang, Willem F. de Boer

## Abstract

Habitat availability determines the distribution of migratory waterfowl along their flyway, which further influences the transmission and spatial spread of avian influenza viruses (AIVs). The extensive habitat loss in the East Asian-Australasian Flyway (EAAF) may have potentially altered the virus transmission and spread, but those consequences are rarely studied. We constructed 6 fall migration networks that differed in their level of habitat loss, wherein an increase in habitat loss resulted in smaller networks with fewer sites. The networks were integrated with an agent-based model and a susceptible-infected-recovered model to simulate waterfowl migration and AIV transmission. We found that extensive habitat loss in the EAAF can 1) relocate the outbreaks northwards responding to the distribution changes of wintering waterfowl geese, 2) increase the outbreak risk in remaining sites due to larger bird congregations, and 3) facilitate AIV transmission among wintering geese. Our modelling output suggested that there was a certain system resilience of migration network to confront the site removal. In addition, the outputs were in line with the predictions from the concept of “migratory escape”, affecting the pattern of infection prevalence in the waterfowl population. Our modelling shed light on the potential consequences of habitat loss in transmitting and spreading AIV at the flyway scale, and suggested the driving mechanisms behind these effects, advocating the importance of nature conservation in changing spatial and temporal patterns of AIV outbreak.

**Author summary:** What are the possible consequences of extensive habitat loss on the transmission and spread of avian influenza viruses (AIVs)? We used a logistic regression model to select the suitable habitats of Greater white-fronted goose in the East Asian-Australasian Flyway and treated these habitats as sites to construct 6 fall migration networks that differed in their level of habitat loss (i.e., site removal). We then simulate geese migration in these networks, and explore the impacts of habitat loss on habitat connectivity and AIV transmission. We found the extensive habitat loss can cause relocation of the outbreaks and increase the outbreak risk and AIV transmission. Our modelling outputs suggested a certain network resilience to confront the site loss, and a “migratory escape” to change the spatial and temporal pattern of infection prevalence in the population. Overall, our study showed that land use changes and habitat loss can affect disease distribution and prevalence, suggested the importance of habitat conservation in changing the spatial and temporal pattern of AIVs transmission and spread.

## Introduction

Emerging zoonotic diseases threaten the health of wildlife, poultry animals and people [1]. To better understand these risks, it is necessary to study the spatial and temporal patterns of pathogen transmission and spread under influence of host and vector distribution and their movements. Among the zoonotic pathogens, avian influenza viruses, AIVs, are well-known for their global circulation and frequent outbreaks in recent decades [2]. This is largely attributed to the seasonal migration and congregation of waterfowl, especially those asymptomatic species [3]. One the one hand, fall migration significantly contributes to the global circulation of the AIV because the migratory population always comprise a considerable amount of immunological naïve (i.e., juvenile) individuals which are relatively more susceptible to AIV [4]. On the other hand, dense congregations facilitate outbreaks by increasing direct virus transmission among individuals [5,6], and by shedding viruses into the environment, thereby increasing environmental transmission [7,8]. Due to the annual migration and congregation of migratory waterfowl, the East Asian-Australasian Flyway (EAAF) has been identified as an area with a high risk for AIV outbreak [5,9]. For instance, on the migratory route of Swan goose *Anser cygnoides*, there were more than 20 outbreaks of highly pathogenic AIV between 2004 and 2017 [10]. Especially the wintering range, such as the Yangtze River floodplain, has become a hotspot for AIV outbreaks [11,12]. Therefore, understanding the distributions of waterfowl species and their congregations, especially in their fall migration and wintering area, can help to identify and predict hotspots of AIV infection [13].

Anthropogenic disturbances such as urban development and land reclamation have caused rapid habitat disappear in the EAAF, especially in the Yangtze River floodplain, one of the regions with extensive habitat loss [14]. As a consequence, migratory waterfowl use alternative habitats for refuelling and wintering [15,16]. For instance, at least 27 waterbird species have changed their distribution for wintering [17], and concentrate in fewer remaining habitats [18,19]. These changes in distribution patterns may influence AIV transmission and spread, which, however, has gained little attention.

Greater white-fronted goose *Anser albifrons* is one of the main hosts of AIV and important vector for transmitting and spreading the virus in the EAAF [20,21]. Since their migration has been well documented by previous satellite tracking studies [22,23], this species is therefore an ideal model species to examine the potential consequences of habitat loss on AIV transmission and spread in the EAAF. The Greater white-fronted goose breeds as north as the Lena Delta in Siberia, and seasonally migrates to their wintering grounds in the Yangtze River floodplain, south Japan and south Korea [24]. Their suitable habitats form a relatively narrow, but long migration corridor, which makes the geese distribution vulnerable to habitat loss [24]. Extensive habitat loss in the wintering range may cause the goose to relocate and concentrate, i.e., increasing in numbers in remaining habitats. Both changes may affect AIV transmission and spread. Thus, we expect that the outbreak risks change spatially and temporally under influence of habitat loss [25], with the remaining habitats having greater risk of AIV outbreak.

In this study, we applied 6 different scenarios of habitat loss to fall migration networks of the Greater white-fronted goose in the EAAF, integrated with an agent-based model (ABM) to simulate the migration of waterfowl, and with a susceptible-infected-recovered model (SIR) to simulate the virus transmission in the population and the virus spread among habitats. We explored the possible consequences of habitat loss on the transmission and spread of AIV. More specifically, we aimed to answer three questions: (1) May habitat loss change the spatial distribution of AIV outbreaks because it changed the distribution of the Greater white-fronted goose? (2) May habitat loss increase the risk for outbreak in remaining habitats? (3) May habitat loss increase virus transmission in a migrating population?

## Results

### Goose distribution and outbreaks relocation

Generally, site removal did not cause drastic changes in the geese and outbreak distributions (Fig 1A-D), until very many sites were removed in the last two scenarios, i.e., the scenarios of removing sites with >10% habitat loss and with >0% habitat loss (Fig 1E and F). The last two scenarios led to shrinking of wintering areas, and confined the migratory geese and the outbreaks to a smaller geographic area. Especially under the extreme scenario, i.e., the scenario of removing sites with >0% habitat loss, the wintering geese relocated northwards from the Yangtze River floodplain to the area above 35.9° N where the last wintering sites were located (ID 86, Fig 1F; also see S5 Fig for geese distribution at each site in each time step), and correspondingly, the outbreaks were confined to that area too.

**Fig 1.**
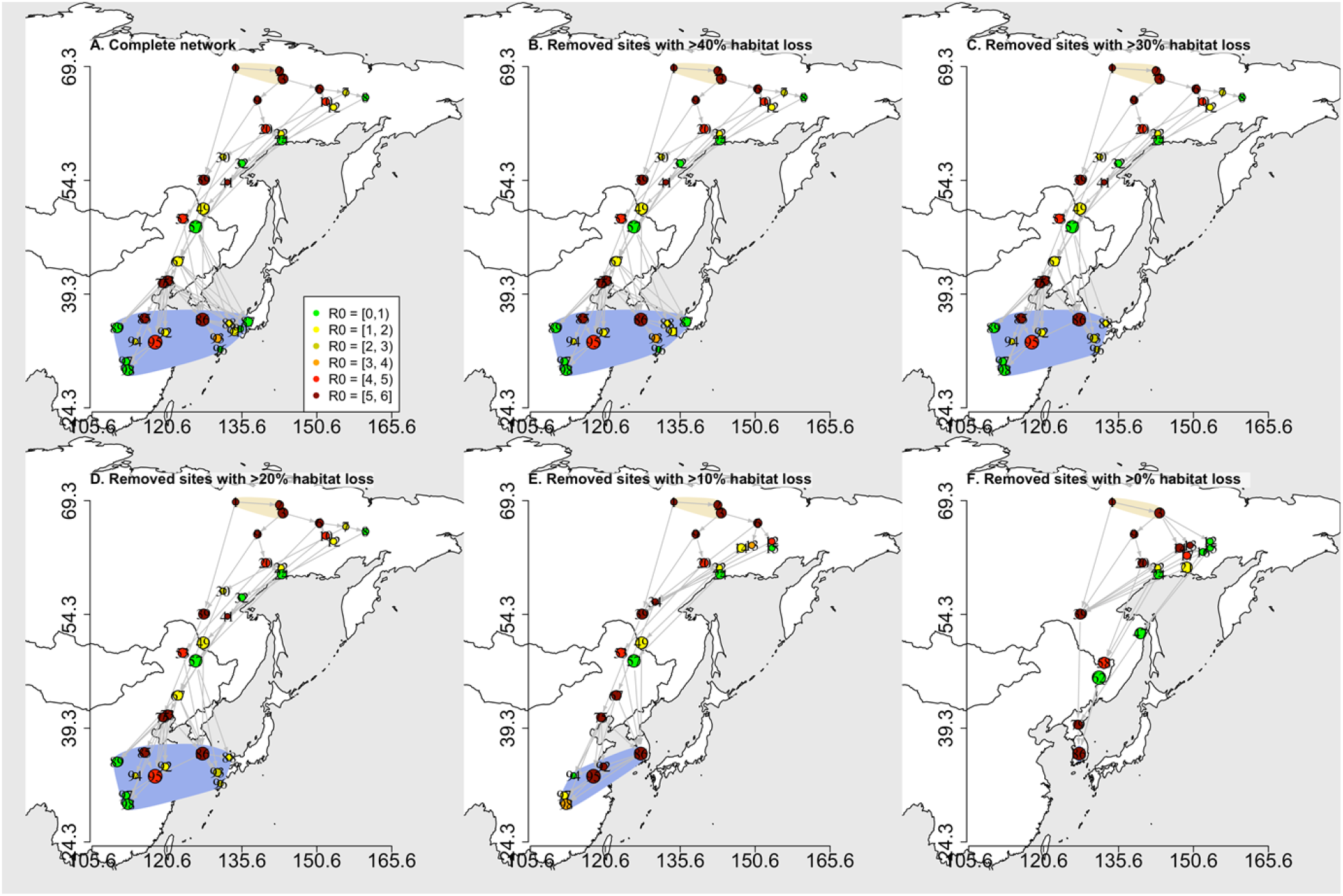
The visited migration networks generated by the simulated geese movement according to the agent-based model. The numbers on the dots are site IDs. The dot colours represent the maximum R_0_ that occurred at the sites during the simulation. The yellow and blue shadows depict the breeding and wintering areas respectively.

In the last two scenarios, in which 16 (16.3%) and 51 (52.0%) sites were removed, respectively (S1 Table), the basic network metrics showed drastic changes compared to the other scenarios (Fig 2). Specifically, the number of visited sites (Fig 2A), the number of sites with outbreak (Fig 2B) and the number of visited links (Fig 2D) declined with increasing site loss (p < 0.001). Whereas the density of visited links (Fig 2E), and the ratio of sites with outbreaks strongly increased (p < 0.001), except for the extreme scenario of which the ratio of sites with outbreak decreased (Fig 2C).

**Fig 2.**
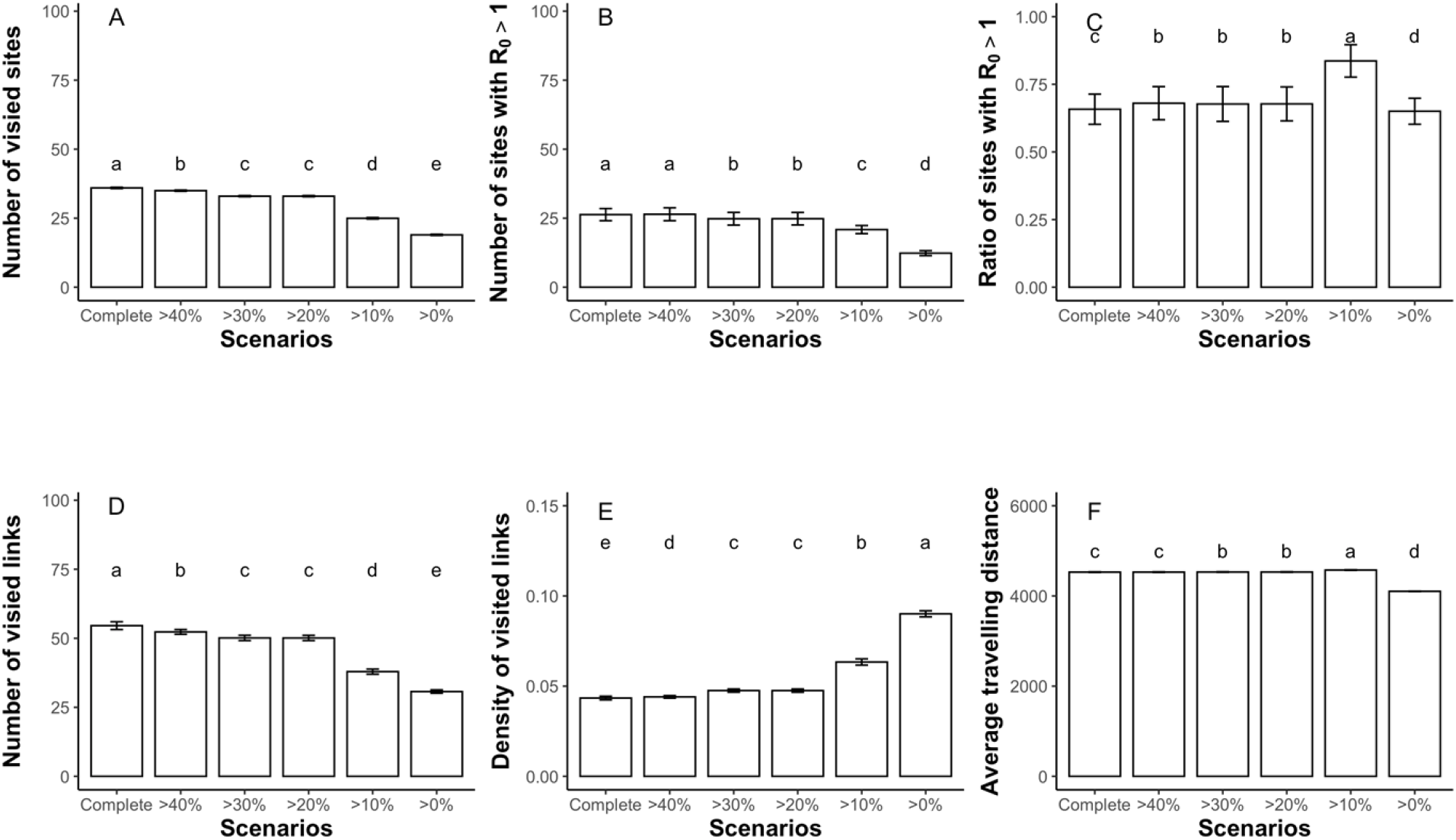
The basic metrics (mean ± SD) of the visited networks under increasing sites removal. A) number of visited sites; B) number of sites with R_0_>1; C) ratio of visited sites with R_0_>1; D) number of visited links; E) density of visited links; F) average travelling distance. The labels on the x axis, from left to right, are the scenario of complete network, and the scenarios of removing sites with >40% habitat loss, with >30% habitat loss, >20% habitat loss, >10% habitat loss and >0% habitat loss. The lowercase annotations indicate the statistic difference of Tukey tests at the level of p=0.001.

The average travelling distance increased from 4,529 (± 6) km in the complete network to 4,575(± 6) km in the scenario of removing sites with >10% habitat loss (Fig 2F; p < 0.001), indicating the geese travelled longer distances due to the site removal. However, it decreased to 4,102 (± 4) km in the scenario of removing sites with >0% habitat loss (Fig 2F; p < 0.001), because the suitable wintering habitats in the Yangtze River floodplain were all removed, causing the southernmost wintering site to relocate northwards (ID 86; Fig 1F). This shortened distance also caused the population to terminate the migration 16 days earlier, compared to that in the complete network (S6 Fig).

### Virus spread over the sites

The GLM showed that the basic reproduction number R_0_ increased with weighted in-degree (Fig 3A and Table 1; p ≤ 0.001), indicating that outbreak risk at site increased under higher connectivity and with more birds migrating from other sites. Moreover, the R_0_ is significantly greater in the last two scenarios, compared to the complete network (Fig 3B and Table 1; p ≤ 0.01), indicating that site removal increased the outbreak risk in the remaining sites.

**Table 1.**
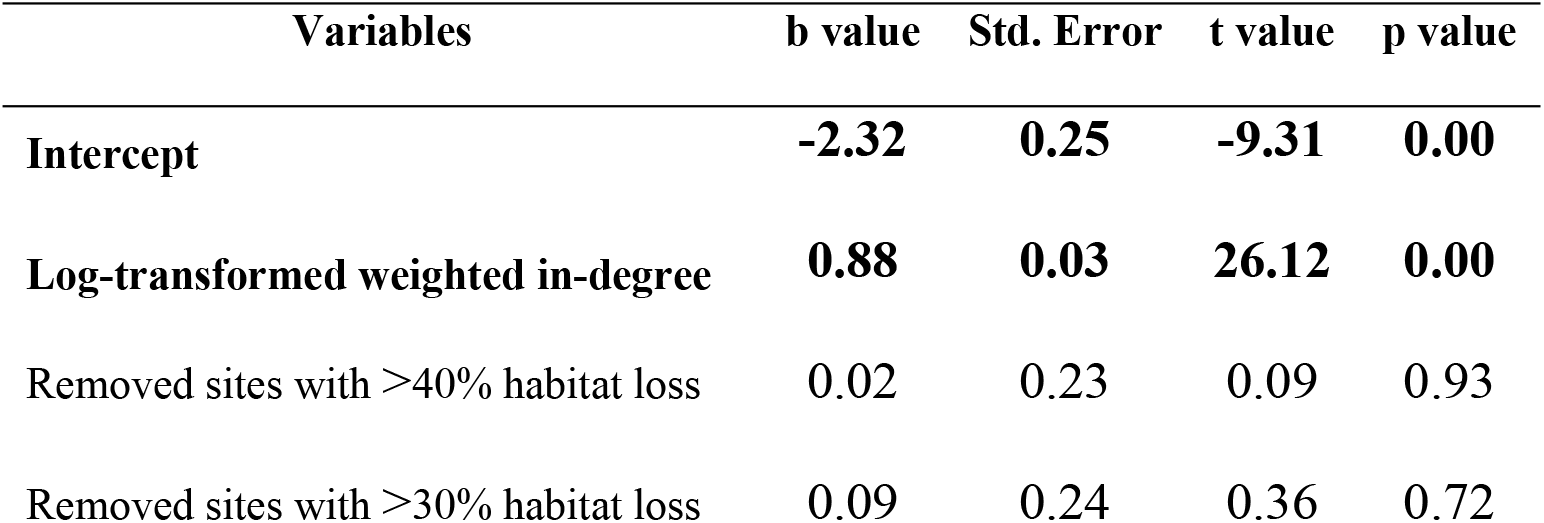
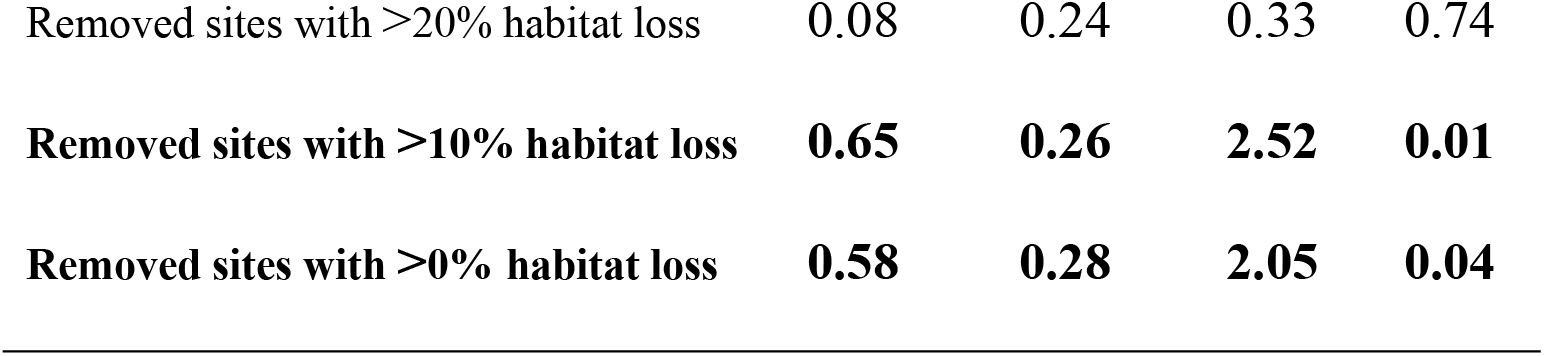
Results of GLM analysis, with coefficients (b), standard error, t-value and p-value. The bold variables were tested significant at the level of p=0.05.

| Variables | b value | Std. Error | t value | p value |
| --- | --- | --- | --- | --- |
| Intercept | **-2.32** | **0.25** | **-9.31** | **0.00** |
| Log-transformed weighted in-degree | **0.88** | **0.03** | **26.12** | **0.00** |
| Removed sites with >40% habitat loss | 0.02 | 0.23 | 0.09 | 0.93 |
| Removed sites with >30% habitat loss | 0.09 | 0.24 | 0.36 | 0.72 |
| Removed sites with >20% habitat loss | 0.08 | 0.24 | 0.33 | 0.74 |
| **Removed sites with >10% habitat loss** | **0.65** | **0.26** | **2.52** | **0.01** |
| **Removed sites with >0% habitat loss** | **0.58** | **0.28** | **2.05** | **0.04** |

**Fig 3.**
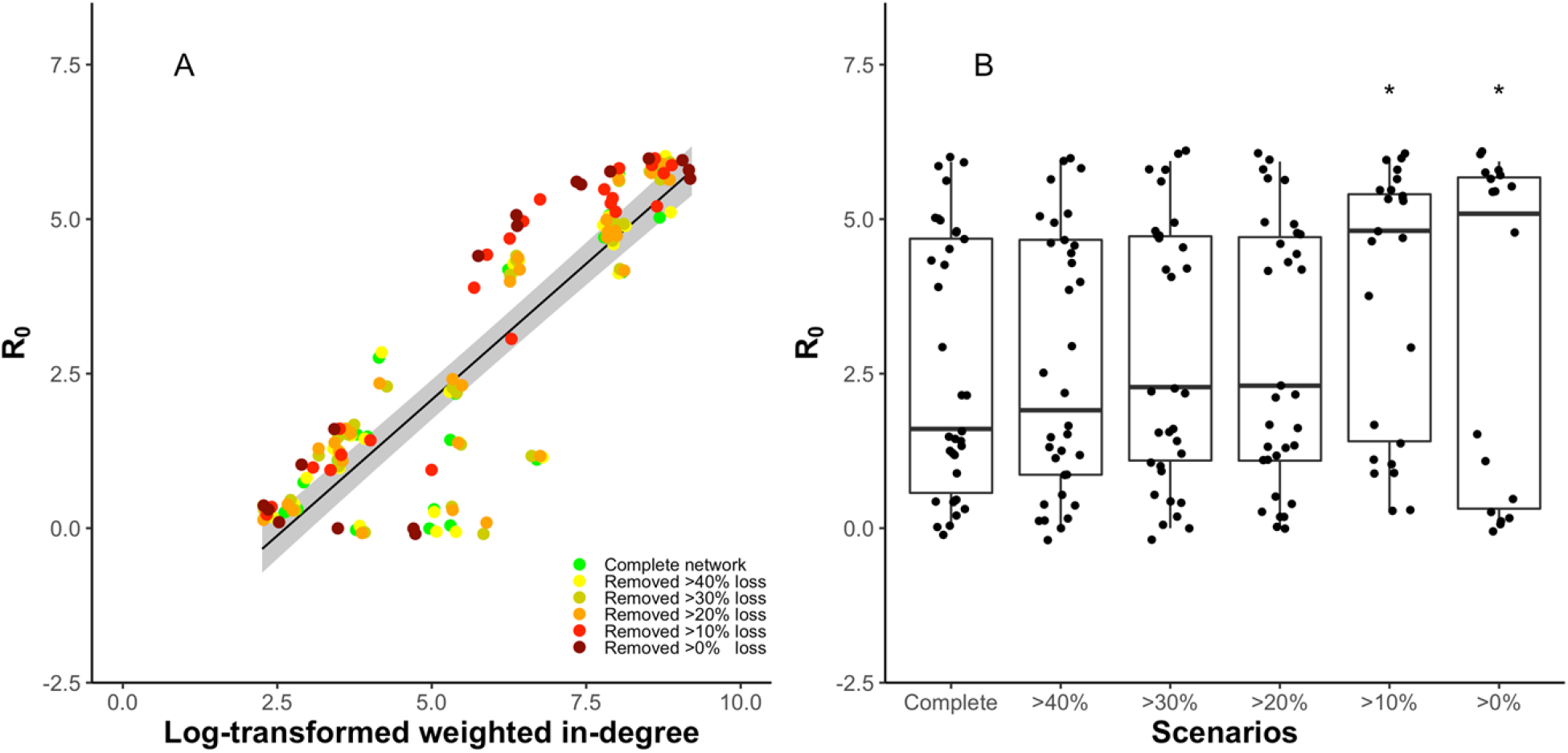
The effects of weighted in-degree and increasing sites loss (scenarios with percentage of site loss) on the basic reproduction number R_0_ at each site. In the panel A, black line represents the GLM fit, and grey shaded area represent the 95% confidence intervals. In the panel B, the labels on the x axis, from left to right, are the scenario of complete network, and the scenarios of removing sites with >40% habitat loss, with >30% habitat loss, >20% habitat loss, >10% habitat loss and >0% habitat loss. The asterisk represents the statistical difference at level of p=0.05, compared to the scenario of complete network. All dots are the R_0_ values generated by agent-based model simulations.

### Virus transmission in the population

The infection prevalence showed similar patterns that one striking infection peak followed by another gentle peak (Fig 4). The first striking peak occurred when the majority geese still congregated at the northmost sites (ID 1; S5 Fig), whereas the second gentle peak coincided with the geese arriving at their wintering sites (Figs S6 and S7) and transmitting the infection (Fig 4B). Moreover, the contribution of the indirect environmental transmission was about 3 times greater than the direct transmission (Fig 4C and D).

**Fig 4.**
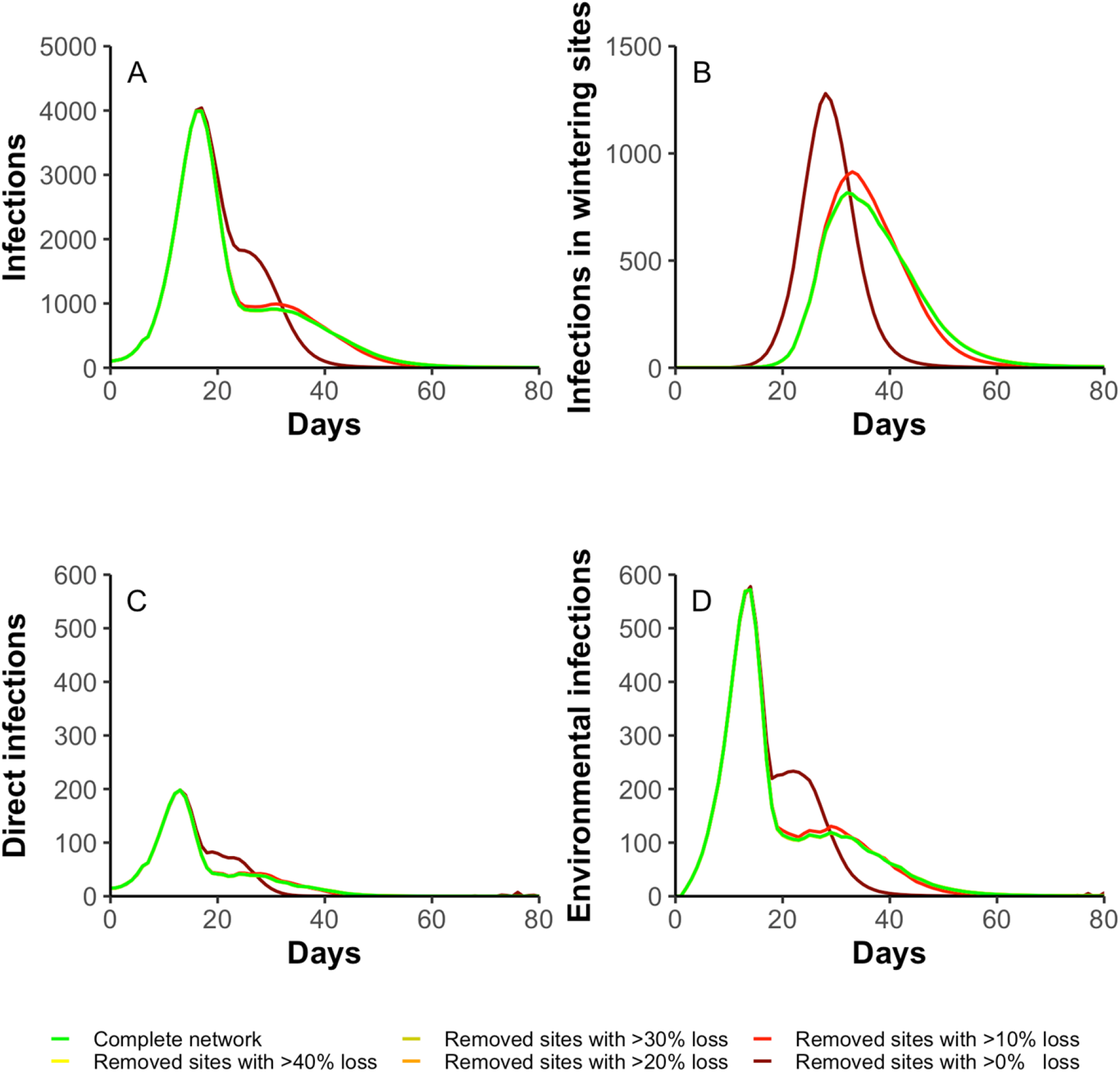
The dynamics of infection prevalence in the migratory population. A) number of infections; B) number of infections in the wintering sites; C) number of infections caused by direct transmission; D) number of infections caused by indirect environmental transmission. The line colours represent the infection dynamics in different scenarios.

The infection prevalence did not differ strongly among the scenarios except for the extreme scenario (Fig 4). Specifically, the second infection peak (i.e., the peak in wintering sites) in the extreme scenario started earlier (Fig 4A) and grew greater (Fig 4B), due to an earlier congregation of geese (Figs S6 and S7). Overall, the infection was spread faster among geese in the extreme scenario.

## Discussion

To gain a better understanding of the effects of anthropogenic disturbance on the spatial and temporal patterns of zoonotic disease outbreaks, we used models to explore the potential consequences of habitat loss on AIV transmission and spread at the flyway scale. By simulating site loss for the Greater white-fronted geese in the EAAF, we found that when migration networks became smaller, with fewer sites, 1) geese and AIV outbreaks relocate northwards, 2) outbreak risk increases among remaining sites, and 3) AIV transmission increases among wintering geese.

The severity of habitat degradation and loss in the Yangtze River floodplain has been calling for attention since 1990s, and this habitat loss is still ongoing [15]. Therefore, in our extreme scenario, all sites located in the Yangtze River floodplain were removed, so that the wintering geese and AIV outbreaks relocated 473 km northwards (Figs 2F and S6). Although such level of site removal is a theoretical approach to explore the potential consequences, the northwards relocation of wintering geese can happen before extensive site disappear, especially for species that are sensitive to habitat degradation [17]. Moreover, other anthropogenic disturbances such as climate change can also contribute to such a relocation [26]. In fact, at least 9 host species have already started moving northwards, abandoning their historical habitats in the Yangtze River floodplain [17].

With removing sites from the networks, some sites turned from low outbreak risk to high risk, e.g., site 13, 34, 58 and 79 (Fig 1 and S2 Table). Especially in the last two scenarios, outbreak risk considerably increased (Fig 3B). Geese movement was the only driver for AIV spreading among sites in our simulation, thus, with more migratory geese arriving at a site from other sites and more connections, the sites had a larger probability to receive infected birds, increasing virus accumulation in the environment (S7 Fig), and triggering an outbreak. In the last two scenarios, the geese had to use fewer remaining sites to finish their migration, and consequently raised the size of congregations and the AIV outbreak risk. This model outcome indicates that larger congregations of waterfowl due to habitat loss might create new AIV hotspots. Field surveys already showed the trend that waterfowl concentrate in fewer remaining habitats. For example, 95% of the Swan goose population, which used to be widely distributed in the Yangtze River floodplain, is nowadays confined to only three major sites [18]. Other species such as the Greylag goose *Anser anser*, Eastern tundra bean goose *Anser serrirostris*, Lesser white-fronted goose *Anser erythropus* and Greater white-fronted goose are showing similar trends, with larger congregations in fewer remaining natural sites [19]. All these changes in spatial distribution and congregation suggested the importance of habitat conservation in affecting outbreak and spread pattern of AIV.

Moreover, the AIV spread was not greatly affected by intermediate levels of habitat loss, because the geese used alternative sites for migration and wintering, until crucial sites for the network connectivity were removed. In other words, removing the sites in order of habitat loss does not immediately alter the spatial distributions of migratory species. However, if the sites that are crucial to the network’s connectivity are not protected or conserved [27], loss of these sites can rapidly change their distributions. Therefore, our simulations of removing sites in order of habitat loss, rather than in the order of decreasing connectivity, might have postponed the more drastic consequences of habitat loss to emerge.

Apart from the spatial changes in AIV outbreaks, extreme site loss also caused changes in the temporal pattern and the prevalence estimates, particularly in the wintering sites. In our simulation, migration behaviour reduced the first infection peak by 35% (2110 geese; S3 Fig), because the migration restricted direct and indirect environmental transmission at the starting site, by allowing susceptible geese to escape from infection [28]. Although these “escaped geese” were eventually infected, longer migration distance can lead to a later congregation and more recoveries, thereby postponing the outbreaks and lowering the maximum prevalence in the wintering sites. However, the extreme scenario led to an earlier geese congregation at wintering sites, the infection was relatively common in the population, thus causing an earlier and larger outbreak. It implies that extreme site removal may exacerbate the AIV outbreak at the end of fall migration, indicating the importance of habitat conservation in potentially controlling AIV outbreaks

The prevalence difference between the sedentary and migratory populations was in line with the predictions from the concept of “migratory escape” [29], i.e., the migration allows host to “escape” from the location where pathogen transmission have accumulated [30]. The migratory escape has been observed in various host-pathogen/parasite relationships, such as warble fly in reindeer [29], or protozoan parasites in butterflies [31]. Although only a few of studies mentioned the migratory escape in the relationship between AIV and migratory waterflow [28,32], it provides new insights to examine the interactions between waterfowl migration and AIV spreading. For example, previous studies found that some waterfowl populations have no AIV infection during their migration, suggesting that their role in spreading AIV was overestimated [4,21,33]. However, this might have been caused by the migratory escape, especially if these samples are taken directly after arrival at their wintering grounds. It indicates that long-distance migration can decrease the prevalence at the end of migration. It also raises the question: what role does migratory escape play in determining the spatial and temporal patterns of AIV infections, especially in a well-preserved migration network? To better answer this question, we call for a larger sampling effort, i.e., not only with a larger sampling period and size [34,35], but also with a larger spatial and species coverage to better capture the spatial and temporal dynamics of AIV infection at flyway scale.

Given our results, we recommend that future studies include population dynamics (e.g., reproduction and age-mediated susceptibility), interaction with other migratory host species, and the incorporation of satellite tracking results to explore the effects of habitat loss. For example, so far we assumed a stable migratory population across all habitat loss scenarios, whereas habitat loss is associated with population declines, especially in the EAAF [18,36,37]. For instance, the Greater white-fronted goose population decreased with 21%-36% from 1990s to 2020 [38]. A declining population might not influence the frequency-dependent direct transmission, but will reduce indirect transmission by decreasing the virus accumulation in the environment [28], which is relatively more important for AIV infection.

In summary, our study explored the potential consequences of habitat loss, i.e., site removal in the model, on spatial and temporal patterns of AIV transmission and spread. Our simulation suggested that there was a certain system resilience, as intermediate sites loss did not greatly affect geese migration patterns, AIV transmission or spread. It is nevertheless interesting to also investigate the impact of migratory escape on AIV transmission and spread. Our study showed that land use changes and habitat loss can affect disease distribution and prevalence. The relationship between conservation, land use changes and pathogen spread can only be well understood if we combine animal tracking studies, disease surveillance, and network analyses.

## Materials and Methods

### Suitable habitats for Greater white-fronted goose

The distribution range of the Greater white-fronted goose in the EAAF covers a large area from 70° N in Russia to 29° N in China, which includes Mongolia, Japan, the Korean Peninsula, and the Yangtze River floodplain. We obtained the classification of breeding region, stopover region, and wintering region on the basis of species distribution maps [39]. Since stopover region and wintering region partly overlap, we classified the wintering region as between 36°N and 29°N.

All potential wetland habitats were extracted from the Global Lakes and Wetlands Database [40], and land cover maps for 1992 and 2012 were obtained from the European Space Agency CCI 300-m annual global land cover products (http://www.esa-landcover-cci.org/). Moreover, the suitability of each potential habitat was estimated by predicting the probability of waterbirds occurrence from a logistic regression model, based on the relationship between the binary observation records, i.e., presence or absence of the Greater white-fronted goose and several environmental predictors, including water body area, elevation, longitude, and suitable foraging areas, i.e., grassland and cropland. The habitat selection followed the procedure described in a previous study [24]. We considered habitats to be suitable when the predicted probability of the goose presence exceeded 75%.

All the suitable habitats were treated as sites in the network construction. We used the coordinate of the geometric centre of each site as the geographic location, and then calculated the geographic distances between all coordinate pairs. The coordinates and distances were calculated with the azimuthal equidistant projection, while the area of each site was calculated with a cylindrical equal area. Moreover, we assigned attributes, including geographic coordinates, area size and type (i.e., breeding, stopover or wintering), to each site, and calculated the percentage of wetland habitat loss between 1992 and 2012. The suitable sites with their attributes are described in the S3 Table.

### Migration network construction

We only generated fall migration networks to test the effect of habitat loss, because the waterfowl migration is more likely to transport the AIV southwards [4,20]. Moreover, simulating a single season and excluding the breeding phase can avoid the complicated infection dynamics that under influence of population dynamics (e.g., changes in population size and age structure). To focus on the effects of habitat loss, we did not differentiate the immunological capacity between adult and juvenile, and excluded the influence of population size changes over time.

We constructed directional links from sites with a higher latitude to sites with a lower latitude between centres of each pair, if the geographical distance *D*_*ij*_ (i.e., geographic distance between site *i* and *j*) was equal or shorter than the migration step length *L*_*step*_ (i.e., the maximum migration distance without rest). In total, we generated 6 theoretical migration networks, by removing the sites with different levels of habitat loss using increments of 10%. These theoretical networks are the complete network, and five network scenarios with increments in habitat loss (i.e., the scenario of removing sites with >40% habitat loss, and >30%, >20%, >10% and >0%). The 6 theoretical networks are shown in S1 Fig, with their corresponding basic network metrics listed in S1 Table.

### Simulation of bird migration

We applied a migratory flow network to simulate the moving geese over the sites [41]. Each site was assigned a variable, site attractiveness 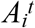 to represent the suitability of the site *i* at given time *t*. Each link was assigned two variables, migration resistance *R*_*ij*_ to represent the difficulty for travelling from site *i* to *j*, and the migration probability *MP*_*ij*_ to represent the likelihood for travelling from site *i* to *j*. Moreover, we assumed the attractiveness *A*_*i*_^*t*^ was negatively influenced by bird density 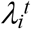, whereas the migration resistance *R*_*ij*_ was positively influenced by geographical distance *D*_*ij*_. These variables at time step *t* were calculated as:

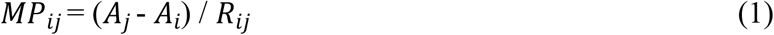

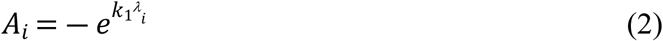

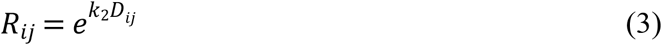

where *k*_*1*_ and *k*_*2*_ are scaling parameters. In general, the decision was determined by bird density and distance between the sites (S1 Appendix), and the bird prefers to select the link with greatest migration probability *MP*_*ij*_.

A total of 10,000 geese were simulated as agents in our model. Each goose was randomly assigned body mass *m*, according to a gaussian distribution at the beginning of simulation. At each time step *t*, the body mass dynamic was calculated as:

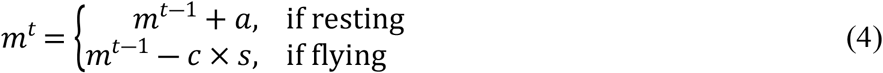

where *a* is the accumulation rate during resting at a site, *c* is the body mass consumption rate during flying, *s* is the flying speed. When a resting bird cumulated enough body mass (i.e., above a threshold *ϕ*), the bird selected a site to migrate to in next time step.

Satellite tracking revealed that Greater white-fronted goose migrates within a narrow corridor (i.e., longitude range) and makes stops for rest and refuelling during fall migration [22,23]. Therefore, we setup two variables, the corridor width *w*, and the expected number of rests *n*, to constrain the sites selection. The corridor width *w* constrains the birds to migrate within a range of longitudes, and the number of rests *n* regulates the number of stopover sites before arriving at the wintering site. The detailed decision-making rules are explained in S1 Appendix. For simplification, we did not include any goal-oriented behaviour or mortality or reproduction in the model.

### Simulation of pathogen transmission

We applied an SIR model to simulate the AIV transmission in the migratory population. A susceptible bird can become infected via either direct transmission, caused by direct contact between susceptible and infected birds, or indirect environmental transmission, caused by viruses in the environment. An infected bird recovered when it has been infected for a certain period *T*_*infection*_. Following previous studies [7,8], we assumed that the birds remained immune after recovery from the infection, as the antibodies to AIV can last for months in Anatidae [42].

Moreover, previous studies found that integrating frequency-dependent transmission and environmental transmission in the model best fitted the observed infection dynamics [7,43]. We therefore followed a previous framework [28], assuming that the direct transmission among birds is frequency-dependent, and the infection probability *ρ* for each susceptible goose at site *i* and time step *t* was calculated as:

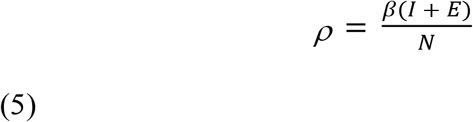

where *β* is the transmission rate parameter, *I* the number of infected geese, *N* the number of geese, *E* the amount of environmental virus at the goose scale (see below).

The amount of virus *V*^*t*^ in the environment at site *i* is calculated as:

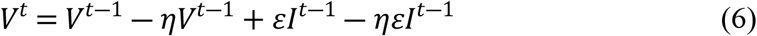

where *η* is the virus decaying rate in the environment, and *ε* is the virus shedding rate. We divided the equation by shedding rate *ε* to obtain:

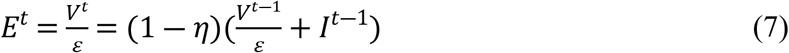

Therefore, we can use the variable *E* to represent the amount of viruses at the scale of goose, for reducing the number of variables [44]. In addition, we only modelled a single AIV strain and one goose population, to avoid the complex infection dynamics caused by cross-immune responses to multiple strains. We further assumed that the infection did not change the migration behaviours of infected geese.

### Model parameterisation

A satellite tracking study suggested that Greater white-fronted goose in the EAAF acquire necessary bodymass stores before starting fall migration [23], and we assume that the resting replenish the energy cost of migration. Therefore, the body mass consumption rate *c* was generally calculated as:

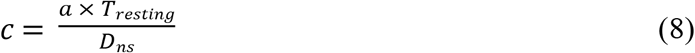

where *T*_*resting*_ is the number of days that Greater white-fronted goose rest on stopover sites, *D*_*ns*_ is the geographic distance between the northernmost and southernmost sites. As the transmission rate parameter *β* of AIV in populations of wild bird is largely unknown [45], we used the value 0.15, which translates to a basic reproduction number R_0_ = 1.03 at the beginning of simulation when no virus existed in the environment, and increasing maximally to 5.97. Other parameters were extracted from published studies (Table 2).

**Table 2.**
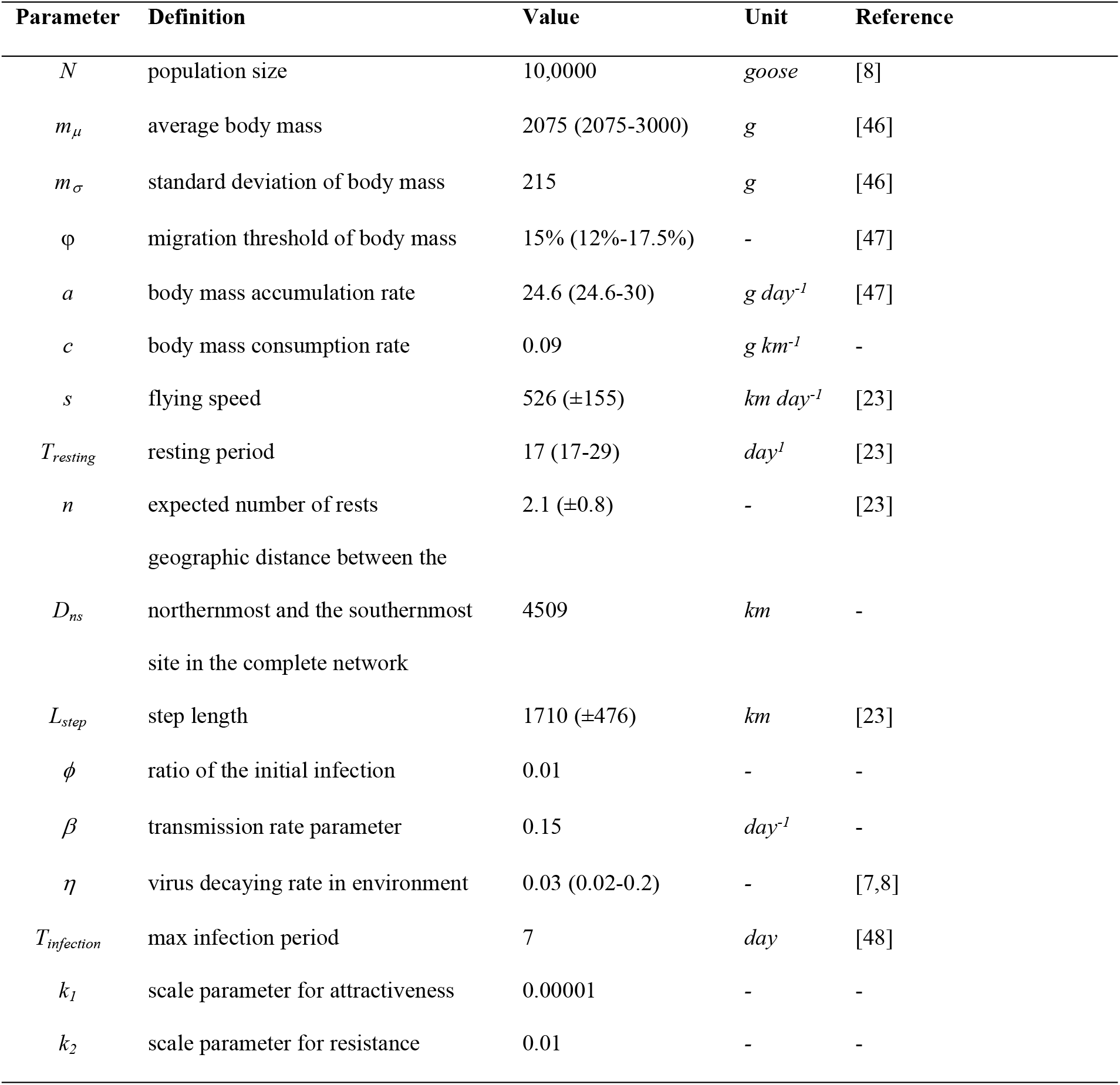
Parameters in the agent-based model, with their abbreviation, definition, value, unit and reference.

### Model analyses

In each network scenario, we initiated the model with all geese at the northernmost site, with an initial infection prevalence 1%. No virus pre-existed in environment at any site at the beginning of the simulations. In the simulation, one time step was equivalent to one day. The simulations ended after all birds stopped migrating and no infected birds existed in the population, which allowed us to capture the full migration and complete prevalence dynamics.

To investigate the infection dynamics during migration, we counted the number of infected birds via direct transmission and environmental transmission. We also calculated the effective reproduction number *R*_*0*_ (i.e., the sum of the average number of new infections) for each site at each time step as [8]:

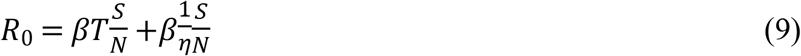

For each network, the simulation iterated 1,000 times and all outputs were averaged. For each iteration, we exported the selected link of each goose at each time step, which were used to generate the visited migration network for carrying out further analysis. Moreover, we used the average link repetitions (i.e., over 1000 iterations) as weight score, and calculated average travelling distance of the population. We used the weighted in-degree of each site to represent both the connection with other sites, and the number of geese arriving on the focal site for resting. Moreover, we also calculated the number of sites, the number of links, and the link densities as network-level metrics to study the effects of habitat removal on network configuration and connectivity [49].

In addition, we conducted a sensitivity analysis for the transmission rate parameter *β* by varying the value by 20% (S2 Fig). To illustrate the effect of migration on the infection dynamics, we also simulated AIV transmission within a sedentary population, with the same parameter setup as the migratory population (S3 Fig). The simulations of the sensitivity analysis and the AIV transmission in the sedentary population were iterated 1000 times, and the outputs were averaged over the iterations, respectively.

### Statistical analyses

We preformed a generalized linear model (GLM) to examine whether the habitat loss can increase the outbreak risk in remaining sites. We selected the maximum *R*_*0*_ across time steps at each site to represent the outbreak risk. The independent predictors were the weighed in-degree (numeric) and network scenarios (categorical). Data generated from all six network scenarios were integrated to perform the GLM analysis. In addition, since the geese were changed their decision-making rules when they were travelling outside and inside of the wintering area (S1 Appendix), we also performed GLM for wintering and other sites (i.e., breeding and stopover sites) separately (S4 Fig and S4 Table). To illustrate the effect of weighted in-degree, we generated the predicted *R*_*0*_ via the GLM by setting the complete network as the reference network scenario. The statistical difference of *R*_*0*_ and the difference of basic network metrics among scenarios were tested via one-way ANOVAs (p<0.05).

In this study, the environmental factors were extracted in ArcMap 10.2.1, the agent-based model was constructed in Netlogo 6.1.1 (S2 Appendix), and all the data processing and statistical analyses were performed in R 4.0.5.

## Acknowledgements

We thank Steve Railsback, Robert Schlicht and Uta Berger for their help in NetLogo programming. We thank Yibin Zheng for checking the code robustness.

## Supporting information captions

**S1 Appendix. Description of the decision-making rules for simulated geese migration**. White boxes represent YES/NO judging procedures; colored boxes represent action procedures; grey boxes are decision-making procedures regarding *liberal migration, rush migration* and *final jump*.

**S2 Appendix. The agent-based model for simulating geese migration and AIV transmission**.

**S1 Fig. The theoretical fall migration networks of Greater white-fronted goose in the EAAF**. The dots are the suitable sites selected by GLM procedure, and the colors represent the level of habitat loss between 1992-2012. The numbers represent site IDs. The yellow and blue shadows are breeding and wintering areas, respectively. The panel A is the complete fall migration network, the other panels are networks with increasing habitat loss with increment of 10%.

**S2 Fig. Infection dynamics generated from the sensitivity analyses by altering the transmission rate parameter *β* with 20% intervals**. Greater *β* increased the size of the first infection peak, but reduced the size of the second peak by quickly consuming susceptible geese in the first infection peak.

**S3 Fig. Comparison of the infection prevalence between sedentary and migratory populations**. The blue lines represent the infection dynamics of the sedentary population, green lines represent infection dynamics of the migratory population under the scenario of a complete network. The migration reduced 35% of the prevalence in the first peak by allowing susceptible geese to escape from infection, but the escaped geese caused the second infection peak when they arrived at their wintering sites.

**S4 Fig. The effects of weighted in-degree and network scenario on the basic reproduction number R**_**0**_ **at each site**. The panels A and C were generated with a GLM of breeding and stopover sites; panels B and D were generated with GLM of wintering sites. In the panel A and B, black lines represent the GLM fit, and grey shaded areas represent the 95% confidence intervals. In the panel C and D, the labels on the x axis, from left to right, are the scenario of complete network, and the scenarios of removing sites with >40% habitat loss, with >30% habitat loss, >20% habitat loss, >10% habitat loss and >0% habitat loss. Asterisk represents the statistical difference at level of p=0.05, compared to the scenario of a complete network. All dots are the R_0_ values generated by agent-based model simulations.

**S5 Fig. The number of geese (log transformed) on each site each day**. The yellow and blue shadows are the breeding and wintering range.

**S6 Fig. The aggregation of geese at wintering sites over time**. The dashed lines mark the time when all geese arrived at the wintering sites. In general, the population finished their migration 16 days earlier under the scenario of removing sites with >0% habitat loss.

**S7 Fig. Dynamics of averaged virus cumulation across all visited sites under each network scenario**.

**S1 Table. Basic network metrics of the 6 theoretical fall migration networks**.

**S2 Table. The visited sites in each scenario, accompanied with the max R**_**0**_ **and number of visiting geese**. The R_0_ is the maximum R_0_ through simulation; the bird is the number of geese that visited the site; the NA indicates the site was removed in the corresponded scenario. Bold records indicate the sites where R_0_ drastically increased (>1).

**S3 Table. Descriptions of the selected suitable sites to generate the migration networks**. The area is the sum of water and grass landscapes (in km^2^) in 1992 and 2012, respectively; the ratio indicates the habitat loss ratio; and the location is the country of the site. B, S, W represent breeding, stopover or wintering sites.

**S4 Table. The regression coefficients (estimate) of separate GLM analysis, with standard errors, t- and p-values**. The bold variables were tested significant at the level of p=0.05.

## Notes

### Competing Interest Statement

The authors have declared no competing interest.

## Reference

1. Jones KE, Patel NG, Levy MA, Storeygard A, Balk D, Gittleman JL, et al. Global trends in emerging infectious diseases. Nature. 2008;451: 990–993. doi:https://doi.org/10.1038/nature06536

2. Verhagen JH, Fouchier RAM, Lewis N. Highly pathogenic avian influenza viruses at the wild–domestic bird interface in Europe: Future directions for research and surveillance. Viruses. 2021;13: 212. doi:https://doi.org/10.3390/v13020212

3. Verhagen JH, Herfst S, Fouchier RAM. How a virus travels the world. Science (80-). 2015;347: 616–617. doi:10.1126/science.aaa6724

4. Kleijn D, Munster VJ, Ebbinge BS, Jonkers DA, Müskens GJDM, Van Randen Y, et al. Dynamics and ecological consequences of avian influenza virus infection in greater white-fronted geese in their winter staging areas. Proc R Soc B Biol Sci. 2010;277: 2041–2048. doi:10.1098/rspb.2010.0026

5. Prosser DJ, Palm EC, Takekawa JY, Zhao D, Xiao X, Li P, et al. Movement analysis of free-grazing domestic ducks in Poyang Lake, China: A disease connection. Int J Geogr Inf Sci. 2016;30: 869–880. doi:10.1080/13658816.2015.1065496

6. van Dijk JG, Verhagen JH, Wille M, Waldenström J. Host and virus ecology as determinants of influenza A virus transmission in wild birds. Current Opinion in Virology. 2018. pp. 26–36. doi:10.1016/j.coviro.2017.10.006

7. Breban R, Drake JM, Stallknecht DE, Rohani P. The role of environmental transmission in recurrent avian influenza epidemics. PLoS Comput Biol. 2009;5. doi:10.1371/journal.pcbi.1000346

8. Rohani P, Breban R, Stallknecht DE, Drake JM. Environmental transmission of low pathogenicity avian influenza viruses and its implications for pathogen invasion. Proc Natl Acad Sci. 2009;106: 10365–10369. doi:10.1073/pnas.0809026106

9. Fearnley L. Wild goose chase: The displacement of influenza research in the fields of Poyang Lake, China. Cult Anthropol. 2015;30: 12–35. doi:10.14506/ca30.1.03

10. Yin S, Xu Y, Batbayar N, Takekawa JY, Si Y, Prosser DJ, et al. Do contrasting patterns of migration movements and disease outbreaks between congeneric waterfowl species reflect differing immunity? Geospat Health. 2021;16. doi:10.4081/gh.2021.909

11. Tian H, Zhou S, Dong L, Van Boeckel TP, Cui Y, Newman SH, et al. Avian influenza H5N1 viral and bird migration networks in Asia. Proc Natl Acad Sci. 2015;112: 172– 177. doi:10.1073/pnas.1405216112

12. Wang D, Li M, Xiong C, Yan Y, Hu J, Hao M, et al. Ecology of avian influenza viruses in migratory birds wintering within the Yangtze River wetlands. Sci Bull. 2021. doi:10.1016/j.scib.2021.03.023

13. Martin V, Zhou X, Marshall E, Jia B, Fusheng G, FrancoDixon MA, et al. Risk-based surveillance for avian influenza control along poultry market chains in South China: The value of social network analysis. Prev Vet Med. 2011. doi:10.1016/j.prevetmed.2011.07.007

14. Mao D, Wang Z, Wu J, Wu B, Zeng Y, Song K, et al. China’s wetlands loss to urban expansion. L Degrad Dev. 2018;29: 2644–2657. doi:https://doi.org/10.1002/ldr.2939

15. Xu Y, Si Y, Wang Y, Zhang Y, Prins HHT, Cao L, et al. Loss of functional connectivity in migration networks induces population decline in migratory birds. Ecol Appl. 2019. doi:10.1002/eap.1960

16. Tellería JL, Pérez-Tris J. Seasonal distribution of a migratory bird: effects of local and regional resource tracking. J Biogeogr. 2003;30: 1583–1591. doi:10.1046/j.1365-2699.2003.00960.x

17. Cao L, Zhang Y, Barter M, Lei G. Anatidae in eastern China during the non-breeding season: Geographical distributions and protection status. Biol Conserv. 2010;143: 650–659. doi:https://doi.org/10.1016/j.biocon.2009.12.001

18. Zhang Y, Cao L, Barter M, Fox AD, Zhao M, Meng F, et al. Changing distribution and abundance of Swan Goose Anser cygnoides in the Yangtze River floodplain: The likely loss of a very important wintering site. Bird Conserv Int. 2011;21: 36–48. doi:10.1017/S0959270910000201

19. Yu H, Wang X, Cao L, Zhang L, Jia Q, Lee H, et al. Are declining populations of wild geese in China ‘prisoners’ of their natural habitats? Current Biology. 2017. pp. R376– R377. doi:10.1016/j.cub.2017.04.037

20. Xu Y, Gong P, Wielstra B, Si Y. Southward autumn migration of waterfowl facilitates cross-continental transmission of the highly pathogenic avian influenza H5N1 virus. Sci Rep. 2016;6: 30262. doi:10.1038/srep30262

21. Yin S, Kleijn D, Müskens GJDM, Fouchier RAM, Verhagen JH, Glazov PM, et al. No evidence that migratory geese disperse avian influenza viruses from breeding to wintering ground. PLoS One. 2017;12: e0177790. doi:10.1371/journal.pone.0177790

22. Kölzsch A, Müskens GJDM, Kruckenberg H, Glazov P, Weinzierl R, Nolet BA, et al. Towards a new understanding of migration timing: slower spring than autumn migration in geese reflects different decision rules for stopover use and departure. Oikos. 2016;125: 1496–1507. doi:10.1111/oik.03121

23. Deng X, Zhao Q, Fang L, Xu Z, Wang X, He H, et al. Spring migration duration exceeds that of autumn migration in Far East Asian Greater White-fronted Geese (Anser albifrons). Avian Res. 2019;10: 1–11. doi:https://doi.org/10.1186/s40657-019-0157-6

24. Xu Y, Si Y, Yin S, Zhang W, Grishchenko M, Prins HHT, et al. Species-dependent effects of habitat degradation in relation to seasonal distribution of migratory waterfowl in the East Asian–Australasian Flyway. Landsc Ecol. 2019;34: 243–257. doi:10.1007/s10980-018-00767-7

25. Gilbert M, Xiao X, Pfeiffer DU, Epprecht M, Boles S, Czarnecki C, et al. Mapping H5N1 highly pathogenic avian influenza risk in Southeast Asia. Proc Natl Acad Sci U S A. 2008;105: 4769–4774. doi:10.1073/pnas.0710581105

26. Crick HQP. The impact of climate change on birds. Ibis (Lond 1859). 2004;146: 48– 56. doi:https://doi.org/10.1111/j.1474-919X.2004.00327.x

27. Xu Y, Si Y, Takekawa J, Liu Q, Prins HHT, Yin S, et al. A network approach to prioritize conservation efforts for migratory birds. Conserv Biol. 2020. doi:10.1111/cobi.13383

28. Yin S, de Knegt HJ, de Jong MCM, Si Y, Prins HHT, Huang ZYX, et al. Effects of migration network configuration and migration synchrony on infection prevalence in geese. J Theor Biol. 2020;502: 110315. doi:10.1016/j.jtbi.2020.110315

29. Altizer S, Bartel R, Han BA. Animal migration and infectious disease risk. Science (80-). 2011;331: 296–302. doi:10.1126/science.1194694

30. Hall RJ, Altizer S, Bartel RA. Greater migratory propensity in hosts lowers pathogen transmission and impacts. J Anim Ecol. 2014;83: 1068–1077. doi:10.1111/1365-2656.12204

31. Satterfield DA, Maerz JC, Altizer S. Loss of migratory behaviour increases infection risk for a butterfly host. Proc R Soc B Biol Sci. 2015;282: 20141734–20141734. doi:10.1098/rspb.2014.1734

32. Arriero E, Muller I, Juvaste R, Martinez FJ, Bertolero A. Variation in immune parameters and disease prevalence among Lesser Black-backed Gulls (Larus fuscus sp.) with different migratory strategies. PLoS One. 2015;10: e0118279. doi:10.1371/journal.pone.0118279

33. Verhagen JH, Van Dijk JGB, Vuong O, Bestebroer T, Lexmond P, Klaassen M, et al. Migratory birds reinforce local circulation of avian influenza viruses. PLoS One. 2014;9. doi:10.1371/journal.pone.0112366

34. Munster VJ, Baas C, Lexmond P, Waldenström J, Wallensten A, Fransson T, et al. Spatial, temporal, and species variation in prevalence of influenza a viruses in wild migratory birds. PLoS Pathog. 2007;3: 0630–0638. doi:10.1371/journal.ppat.0030061

35. Wallensten A, Munster VJ, Latorre-Margalef N, Brytting M, Elmberg J, Fouchier RAM, et al. Surveillance of influenza A virus in migratory waterfowl in northern Europe. Emerg Infect Dis. 2007;13: 404–411. doi:10.3201/eid1303.061130

36. Studds CE, Kendall BE, Murray NJ, Wilson HB, Rogers DI, Clemens RS, et al. Rapid population decline in migratory shorebirds relying on Yellow Sea tidal mudflats as stopover sites. Nat Commun. 2017. doi:10.1038/ncomms14895

37. Chen S, Zhang Y, Borzée A, Liang T, Zhang M, Shi H, et al. Landscape attributes best explain the population trend of wintering greater white-fronted goose (Anser albifrons) in the Yangtze River Floodplain. Land. 2021;10: 865. doi:https://doi.org/10.3390/land10080865

38. Deng X, Zhao Q, Solovyeva D, Lee H, Bysykatova-Harmey I, Xu Z, et al. Contrasting trends in two East Asian populations of the Greater White-fronted Goose Anser albifrons. Wildfowl. 2020; 181–205.

39. Birdlife International and NatureServe. Bird species distribution maps of the world Version 5.0. 2015 [cited 2 Nov 2016]. Available: http://www.birdlife.org

40. Lehner B, Döll P. Development and validation of a global database of lakes, reservoirs and wetlands. J Hydrol. 2004;296: 1–22. doi:https://doi.org/10.1016/j.jhydrol.2004.03.028

41. Taylor CM, Laughlin AJ, Hall RJ. The response of migratory populations to phenological change: A Migratory Flow Network modelling approach. J Anim Ecol. 2016;85: 648–659. doi:10.1111/1365-2656.12494

42. Samuel MD, Hall JS, Brown JD, Goldberg DR, Ip H, Baranyuk V V. The dynamics of avian influenza in Lesser Snow Geese: Implications for annual and migratory infection patterns. Ecol Appl. 2015;25: 1851–1859. doi:10.1890/14-1820.1

43. Roche B, Lebarbenchon C, Gauthier-Clerc M, Chang CM, Thomas F, Renaud F, et al. Water-borne transmission drives avian influenza dynamics in wild birds: The case of the 2005-2006 epidemics in the Camargue area. Infect Genet Evol. 2009;9: 800–805. doi:10.1016/j.meegid.2009.04.009

44. Bravo De Rueda C, De Jong MC, Eblé PL, Dekker A. Quantification of transmission of foot-and-mouth disease virus caused by an environment contaminated with secretions and excretions from infected calves. Vet Res. 2015;46. doi:10.1186/s13567-015-0156-5

45. Lisovski S, van Dijk JGB, Klinkenberg D, Nolet BA, Fouchier RAM, Klaassen M. The roles of migratory and resident birds in local avian influenza infection dynamics. J Appl Ecol. 2018; 1–13. doi:10.1111/1365-2664.13154

46. Dunning Jr. JB. CRC Handbook of Avian Body Masses. CRC handbook of avian body masses. Second ed. 2008. doi:10.1017/S0963180113000479

47. Fox AD. The Greenland white-fronted goose Anser albifrons flavirostris: The annual cycle of a migratory herbivore on the European continental fringe. National Environmental Research Institute, Denmark. 2003. Available from: www.dmu.dk

48. Hénaux V, Samuel MD. Avian influenza shedding patterns in waterfowl: Implication for surveillance, environmental transmission, and disease spread. J Wildl Dis. 2011;47: 566–578. doi:10.7589/0090-3558-47.3.566

49. Banks NC, Paini DR, Bayliss KL, Hodda M. The role of global trade and transport network topology in the human-mediated dispersal of alien species. Ecol Lett. 2015;18: 188–99. doi:10.1111/ele.12397

